# *daf-16/FOXO* activity before dauer diapause anticipates dauer exit and post-dauer development via DAF-12/NHR

**DOI:** 10.1101/2025.06.18.660181

**Authors:** Matthew J. Wirick, Isaac T. Smith, Benjamin S. Olson, Amelia F. Alessi, Himani Galagali, Kyal Lalk, Mikayla N. Schmidt, Kevin J. Ranke, Lane S. Rinkus, Xantha Karp

## Abstract

Animal development occurs in the context of varying environmental conditions. Many animal species can withstand adverse conditions by entering a stress-resistant and developmentally arrested diapause stage. If conditions improve, animals can recover and complete development. In *Caenorhabditis elegans* larvae that encounter adverse environments, dauer diapause can interrupt developmental progression after the second larval molt. During continuous (non-dauer) development, a heterochronic molecular timer comprised primarily of microRNAs and their targets controls the progression of stage-specific cell fates in lateral hypodermal seam cells. In unfavorable conditions, the DAF-16/FOXO transcription factor promotes dauer formation in part through opposing production of the ligand for the DAF-12 nuclear hormone receptor. Ligand-free DAF-12 promotes dauer formation and represses expression of heterochronic microRNAs in the *let-7* family. Here, we show that *daf-16* acts before dauer formation in anticipation of growth after dauer by promoting both dauer exit and developmental progression. We found that *daf-16(0)* post-dauer adults showed delayed dauer exit, reiterative heterochronic defects, and reduced expression of *let-7-*family microRNAs. These phenotypes were suppressed by addition of the DAF-12 ligand, dafachronic acid. Dafachronic acid is synthesized from cholesterol, and dauer larvae sequester cholesterol in the intestinal lumen until dauer exit. We found that a fluorescent cholesterol analog was not retained in *daf-16* mutant larvae during dauer recovery. Timed auxin-mediated depletion of DAF-16 indicated that *daf-16* is required before dauer formation to prevent reiterative seam cell fates in post-dauer animals. We propose a model whereby *daf-16* acts prior to dauer formation to enable dafachronic acid synthesis by retaining cholesterol during dauer recovery. Ligand-bound DAF-12 then promotes dauer exit and expression of *let-7*-family microRNAs, thereby promoting developmental progression. Thus, *daf-16* coordinates dauer entry with anticipation of recovery and post-dauer development.

## Results and Discussion

### *daf-16* promotes post-dauer adult cell fate

Depending on environmental conditions, *C. elegans* larvae can either develop continuously through four larval stages (L1-L4) before reaching adulthood, or through a life history that includes dauer diapause (Figure 1A). Developmental progression in the dauer life history differs from continuous development at several steps, including a lengthened pre-dauer L2d stage, the suspension of development during dauer diapause, and, if conditions improve, the resumption of development after a recovery process^1^. Dauer formation is regulated by three interconnected signaling pathways: insulin/IGF signaling (IIS), TGFβ, and the DAF-12 nuclear hormone receptor (NHR) signaling (Figure 1B)^2^. The DAF-16/FOXO transcription factor acts downstream of IIS to promote dauer diapause in adverse environments^3,4^. We previously showed that *daf-16* contributes to developmental arrest during dauer by blocking the precocious adoption of adult cell fate in the lateral hypodermis, including lateral hypodermal seam cells^5,6^. Here we show that *daf-16* also regulates post-dauer seam cell fate and dauer exit through a distinct mechanism involving DAF-12/NHR, as described in the sections below.

**Figure 1.**
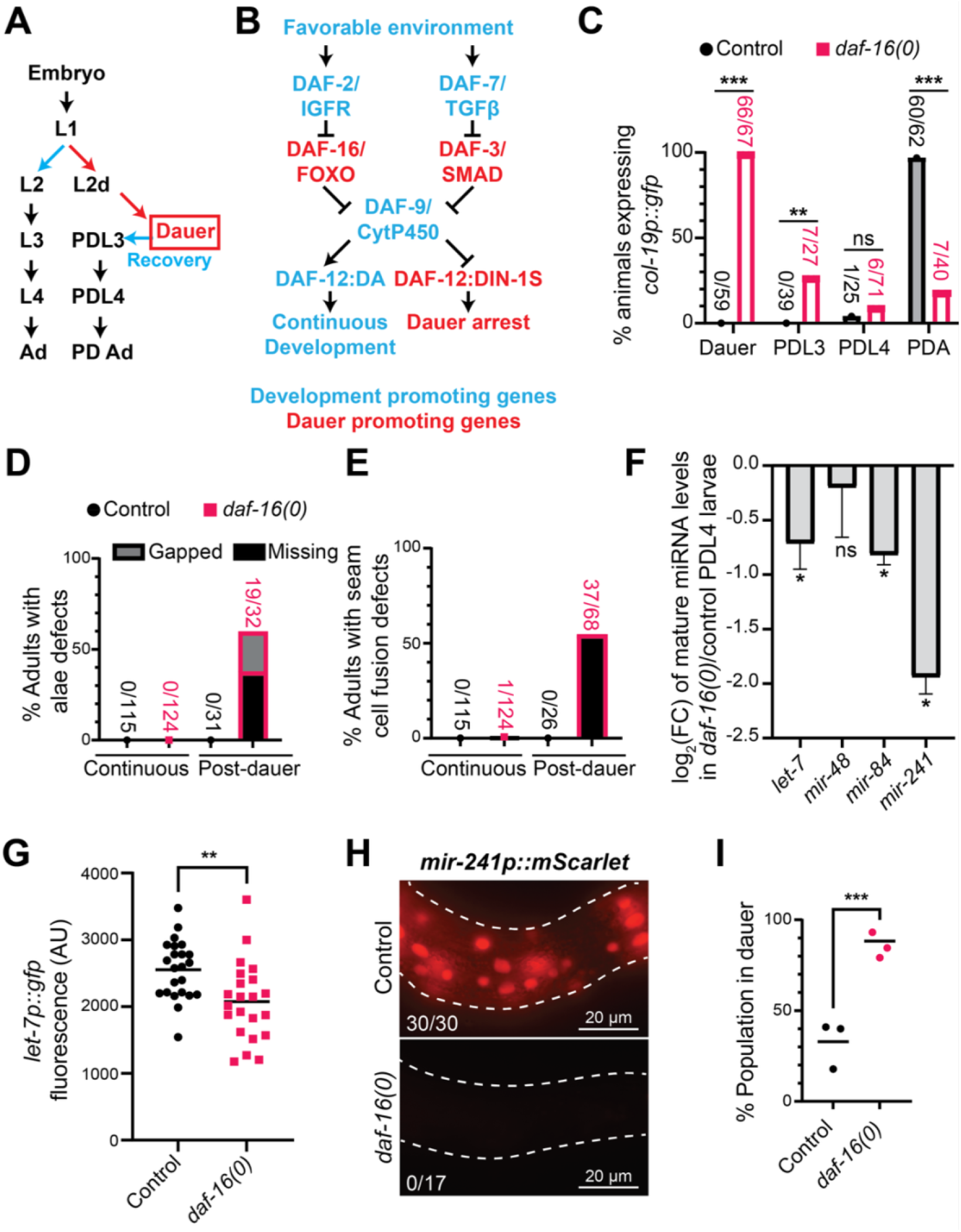
*daf-16* coordinates the exit from dauer and the resumption of seam cell development after dauer. (A) Continuous (left) and dauer (right) life histories in *C. elegans.* Favorable environmental conditions (blue arrow) sensed in the L1 stage promote continuous development, beginning with entry into the L2 stage and culminating in adulthood (Ad). Adverse conditions (red arrow) induce L1 stage animals to molt into the pre-dauer L2d stage. Continued adverse conditions promote entry into dauer diapause. If dauer larvae encounter favorable environments, they will exit dauer and begin a recovery process. After recovery, dauer larvae re-enter development through post-dauer (PD) larval stages. (B) Genetic model of the dauer decision pathways. (C) The penetrance of *daf-16(0)* animals expressing *col-19p::gfp* is reduced after dauer and remains low in all PD stages, including in adults. The dauer penetrance data come from Wirick et al., 2021^5^. **p=0.0011, ***p<0.001, 2X2 Fisher’s exact test, two tailed. (D) *daf-16(0)* PD adults had defects in adult alae synthesis. In C and subsequent figures showing adult alae, numbers indicate the number of adults with defects, defined as missing or gapped alae, over the total number of adults assessed. (E) *daf-16(0)* PD adults showed defects in seam cell fusion, defined as animals expressing the apical junction marker *ajm-1::gfp* between at least one pair of adjacent seam cells. (F) *daf-16(0)* PDL4 larvae expressed reduced levels of mature *let-7* family microRNAs. miRNA levels were normalized to U18. *p<0.05, t-test, two-tailed. (G) *daf-16(0)* PDL4 larvae expressed reduced levels of *let-7p::gfp*. Each data point represents the expression of *let-7p::gfp* throughout the body of one PDL4 larva. **p=0.0015, Mann-Whitney test, two-tailed. (H) *daf-16(0)* larvae at the PDL3 molt (PDL3m) showed greatly reduced expression of an endogenous *mir-241* transcriptional reporter. Numbers indicate the number of PDL3m larvae with robust *mir-241p::mScarlet* expression (see Figure S3) over the total number of larvae assessed. Dashed lines show the outline of the worm. (I) *daf-16(0)* larvae show delayed dauer exit. Each data point represents the population of larvae remaining in dauer 24 hours after shifting dauer larvae from 24°C to 20°C in an individual experiment (n=20-29 larvae per data point). ***p<0.001, 2X2 Fisher’s exact test, one-tailed.

To determine the role for *daf-16* in dauer exit and post-dauer development, we created strains with the *daf-16(mgDf50)* null allele (hereafter *daf-16(0))* and the *daf-7(e1372)* allele which causes dauer entry at 24°C or 25°C. The *daf-7* allele was used to induce dauer formation in the dauer-defective *daf-16(0)* background^3,7^. This allele was present in the background of all strains that developed through the dauer life history, and for simplicity is not always mentioned. (See Table S1 for a full list of genotypes). To induce dauer exit, dauer larvae were picked to new plates and shifted to 15°C or 20°C.

During dauer, *daf-16(0)* mutants precociously express the *col-19p::gfp* adult cell fate marker^5^. We asked if the expression *of col-19p::gfp* persisted as *daf-16(0)* larvae exited dauer and resumed development as post-dauer (PD) animals. We found that *col-19p::gfp* expression declined over time and, surprisingly, this adult cell fate marker remained largely off in PD adults (Figure 1C). Furthermore, *daf-16(0)* PD adults showed penetrant defects in two additional adult-specific characteristics: the expression of adult alae, a cuticular structure secreted by seam cells upon the larval-to-adult switch, and the loss of adherens junctions that are present between larval but not adult seam cells (Figure 1D-E). These phenotypes were only observed in PD adults; *daf-16(0)* adults that developed through the continuous life history expressed adult characteristics similar to wild-type controls (Figure 1 D-E)^5^. Consistent with the reduced expression of these adult characteristics, we found that *daf-16(0)* PD adults failed to downregulate an L4 hypodermal cell fate marker, *col-38p::yfp* (Figure S1)^8^. Taken together, these results indicate that *daf-16* functions specifically in the dauer life history to promote adult seam cell fate.

### *daf-16* promotes transcription of *let-7-*family microRNAs

Mutations of heterochronic microRNAs (miRNAs) result in reiterative phenotypes, similar to those we saw in *daf-16(0)* PD adults^9–11^. To broadly test if *daf-16* regulates miRNAs to control the larval-to-adult switch, we crossed worms with a null mutation in one of the major miRNA-specific Argonaute proteins, *alg-1(0)*, into our *daf-16(0)* background. *alg-1(0)* mutants display reiterative phenotypes during continuous development but have few defects after dauer, creating a miRNA-sensitized background to study post-dauer adult cell fate^12,13^. Indeed, when observing alae in PD adults, we found an enhancement in alae defects in the *daf-16(0); alg-1(0)* double mutant relative to the *daf-16(0)* single mutant (Figure S2), suggesting that *daf-16* functions with miRNAs to regulate adult cell fate after dauer.

During dauer, *daf-16* opposes expression of *let-7-*family miRNAs which indirectly promote adult cell fate^6^. There are four *let-7-*family miRNAs that act in the heterochronic pathway: *mir-48, mir-84, mir-241,* and *let-7*^9,11^. To test whether *daf-16* regulates the *let-7* family miRNAs after dauer, we used qRT-PCR to quantify the levels of mature *let-7*-family miRNAs in *daf-16(0)* and control PDL4 larvae and found that *mir-84*, *mir-241*, and *let-7* were downregulated in *daf-16(0)* PD larvae (Figure 1F). This downregulation appears to be mediated transcriptionally because two transcriptional reporters, *let-7p::gfp* and *mir-241p::mScarlet,* were both downregulated in *daf-16(0)* PD larvae (Figure 1G-H, Figure S3).

### *daf-16* is required for timely exit from dauer diapause

In the course of performing the experiments described above, we noticed that *daf-16(0)* mutants required an extra day to recover from dauer, relative to control larvae. This dauer exit phenotype was initially surprising, since *daf-16* activity is required to maintain dauer arrest^14,15^. However, delayed dauer exit in *daf-16* mutants was also observed in a prior report^16^. To quantify this phenotype in our hands, we measured the percentage of dauer larvae that failed to recover 24 hours after shifting dauer larvae from 24°C to 20°C and observed that the majority of *daf-16(0)* larvae remained in dauer, while control larvae had largely resumed development at this time (Figure 1I). This led us to ask if the additional time that *daf-16(0)* larvae spent in dauer contributed to the adult cell fate defects. We assessed alae formation in control and *daf-16(0)* populations at 2 and 3 days after shifting dauer larvae from 24°C to 20°C. At 2 days, most control animals and a few *daf-16(0)* animals had reached adulthood. At 3 days, most *daf-16(0)* animals had reached adulthood and some of the control animals that had not reached adulthood at 2 days were now adults. The control PD adults had very few alae defects at either 2 or 3 days, indicating that time in dauer *per se* does not lead to alae defects (Figure S4). However, *daf-16(0)* 2-day post-recovery adults had substantially fewer alae defects than the *daf-16(0)* 3-day post-recovery adults (Figure S4). These results suggest that for *daf-16* mutants, the dauer exit and adult cell fate phenotypes may be connected. Experiments described in the sections below are consistent with this interpretation.

### *daf-16* promotes the liganded DAF-12 complex

Thus far we have shown that *daf-16* promotes dauer exit, PD expression of the *let-7* family miRNAs, and PD adult cell fate. The regulation of this suite of processes is reminiscent of the role of the DAF-12 nuclear hormone receptor. In favorable conditions, the DAF-12 ligand, dafachronic acid (DA) is produced, and the DAF-12:DA complex directly activates *let-7-*family miRNA transcription and promotes continuous development. In adverse conditions, ligand-free DAF-12 binds its corepressor DIN-1S to repress *let-7-*family miRNA transcription and promotes dauer entry and dauer maintenance (Figure 1B)^17–20^. Furthermore, if dauer larvae find favorable environments, the production of DA is required for dauer exit^21^. We hypothesized that *daf-16(0)* PD animals may have elevated activity of the repressive DAF-12 complex, leading to delayed dauer exit and reduced levels of *let-7-*family miRNAs. For technical reasons, we were unable to test *daf-12* loss-of-function mutants (see Figure S5). However, we reasoned that if our hypothesis was correct, boosting the DAF-12:DA complex should suppress the *daf-16(0)* PD defects.

To shift the repressive ligand-free DAF-12 complex to the activating DAF-12:DA complex, we treated *daf-16(0)* dauer larvae with 100nM Δ7-DA and assessed timing of dauer exit and alae morphology in PD adults. Treatment with Δ7-DA almost completely suppressed alae defects in *daf-16(0)* PD adults and restored timely dauer exit, while *daf-16(0)* animals treated with ethanol as a carrier control retained penetrant alae and dauer exit defects (Figure 2A-C). We also tested intermediates in the DA biosynthetic pathway (Figure 2A) to determine if they may also suppress these phenotypes. All intermediates tested suppressed alae and dauer exit defects in *daf-16(0)* animals to some degree, except for the starting sterol, cholesterol, which is already in abundance in the NGM media (Figure 2B-C). The suppression of adult alae defects in *daf-16(0)* PD adults by Δ7-DA treatment coincided with the rescue of *let-7p::gfp* and *mir-241p::mScarlet* expression in PD larvae (Figure 2D-E and Figure S6). Addition of Δ7-DA had no effect on levels of these transcriptional reporters in the control *daf-16(+)* background, suggesting that DA is limiting in the *daf-16(0)* background but not in controls (Figure 2D-E and Figure S6).

**Figure 2.**
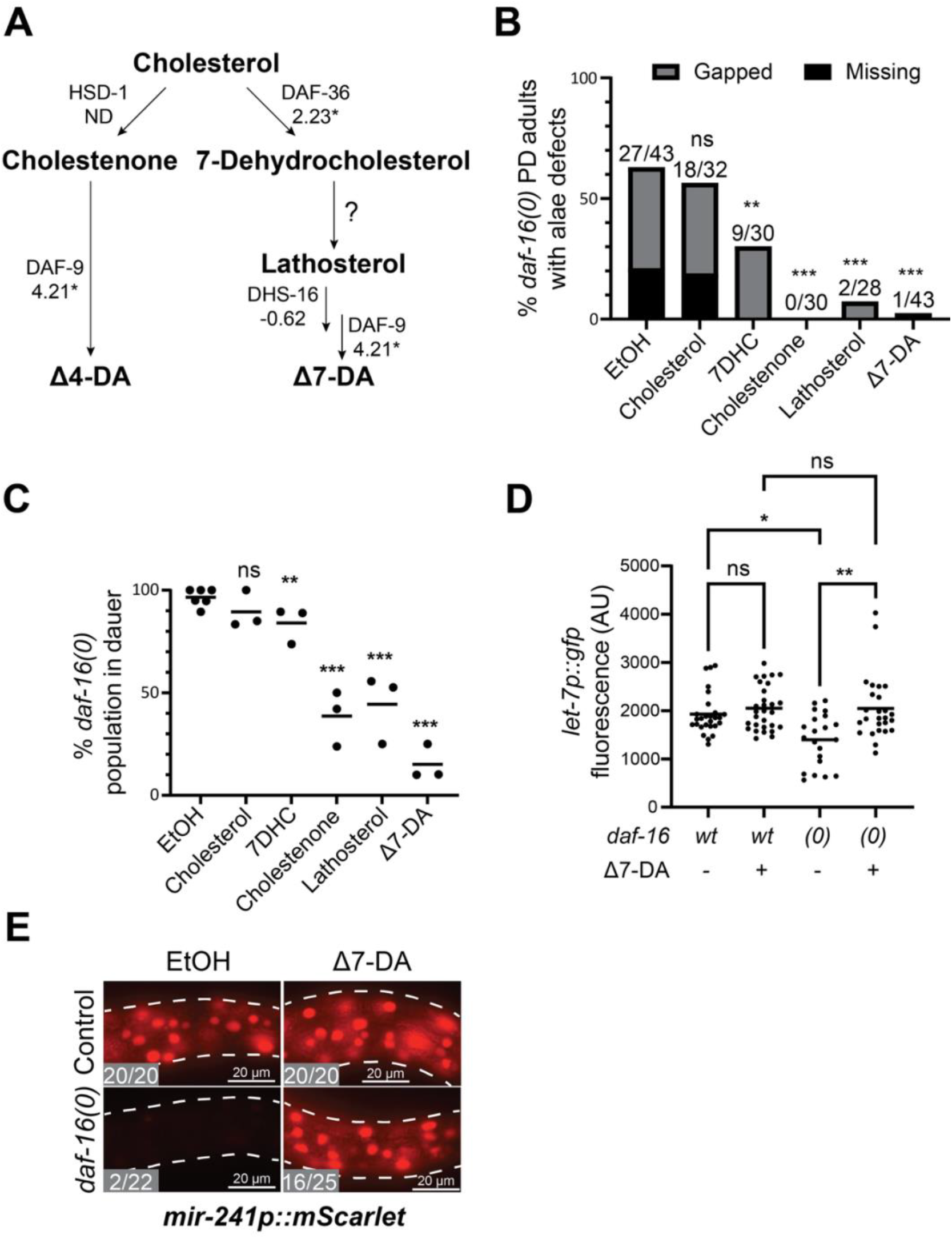
*daf-16* promotes the production of dafachronic acid to regulate adult cell fate. (A) Model of the biosynthetic pathway for the two originally described forms of DA, Δ4-DA and Δ7-DA^35,36^. Chemical compounds are highlighted in bold and the enzymes that catalyze each reaction are listed. Numbers below enzyme names indicate the log_2_(fold change) in expression of each gene in *daf-16(0); daf-7* vs *daf-7* dauer larvae, from Wirick et al., 2021^5^. Asterisks indicate that the change in gene expression was statistically significant (FDR < 0.05). ND = Not detected in either sample. (B) Treatment of 100nM Δ7-DA or 33.3µM of intermediates downstream of cholesterol suppressed alae defects in *daf-16(0)* PD adults, while treatment with 33.3µM cholesterol appeared similar to the control EtOH carrier. **p=0.003, ***p<0.001, 2X3 Fisher’s exact test, two-tailed. (C) Treatment of 100nM Δ7-DA or 33.3µM of intermediates downstream of cholesterol suppressed dauer recovery defects in *daf-16(0)* PD adults, whereas 33.3µM cholesterol did not (n=16-21 larvae per data point). **p=0.006, ***p<0.001, 2X2 Fisher’s exact test, one-tailed. Each data point represents the population of animals that remained in dauer 24 hours after shifting dauer larvae from 24°C to 20°C in an individual experiment. (D) Treatment of 100nM Δ7-DA restored *let-7p::gfp* expression in *daf-16(0)* PDL4 larvae. Each data point represents the whole-body expression of *let-7p::gfp* for one PDL4 larva. *p=0.029, **p<0.01, Kruskal-Wallis test with Dunn’s multiple comparisons test. (E) Treatment of Δ7-DA restored robust *mir-241p::mScarlet* expression in *daf-16(0)* PDL3m larvae. Numbers indicate the number of PDL3m larvae with robust *mir-241p::mScarlet* expression (see Figure S6) over total number of larvae assessed. Dashed lines outline worm cuticles.

### *daf-16* promotes the expression of *scl-12* and *scl-13*

We next asked what the mechanism by which *daf-16* promotes the liganded DAF-12 complex might be. A simple hypothesis is that *daf-16* might promote the expression of an enzyme within the DA biosynthetic pathway. However, mRNA-seq data comparing *daf-16(0)* to control dauer larvae did not show downregulation of known enzymes in this pathway (Figure 2A)^5^. Furthermore, as described above, multiple intermediates in the DA biosynthetic pathway suppressed PD *daf-16(0)* defects (Figure 2B-C). These results suggest that DA may be limiting in *daf-16(0)* PD larvae and that *daf-16* promotes DA production somewhere upstream of the conversion of cholesterol to 7-dehydrocholesterol (Figure 2A).

Prior work demonstrated that cholesterol is sequestered in the intestinal lumen of dauer larvae, and that this cholesterol sequestration is required to synthesize DA upon dauer recovery^22^. Little is known about this process, but it appears to be mediated by *scl-12* and *scl-13* which encode sterol-binding proteins^22^. To ask if *daf-16* might regulate the expression of these genes, we examined our prior mRNA-sequencing data from *daf-16(0)* and control dauer larvae^5^ and found that both *scl-12* and *scl-13* are strongly downregulated in *daf-16(0)* dauer larvae (Figure 3A). Furthermore, our prior ChIP-seq data from starved dauer larvae with endogenous DAF-16 tagged with 3xFLAG^6^ showed DAF-16 enrichment in two peaks in the shared promoter region between *scl-12* and *scl-13,* suggesting that DAF-16 may directly bind the promoters of these genes to activate their expression (Figure 3B).

**Figure 3.**
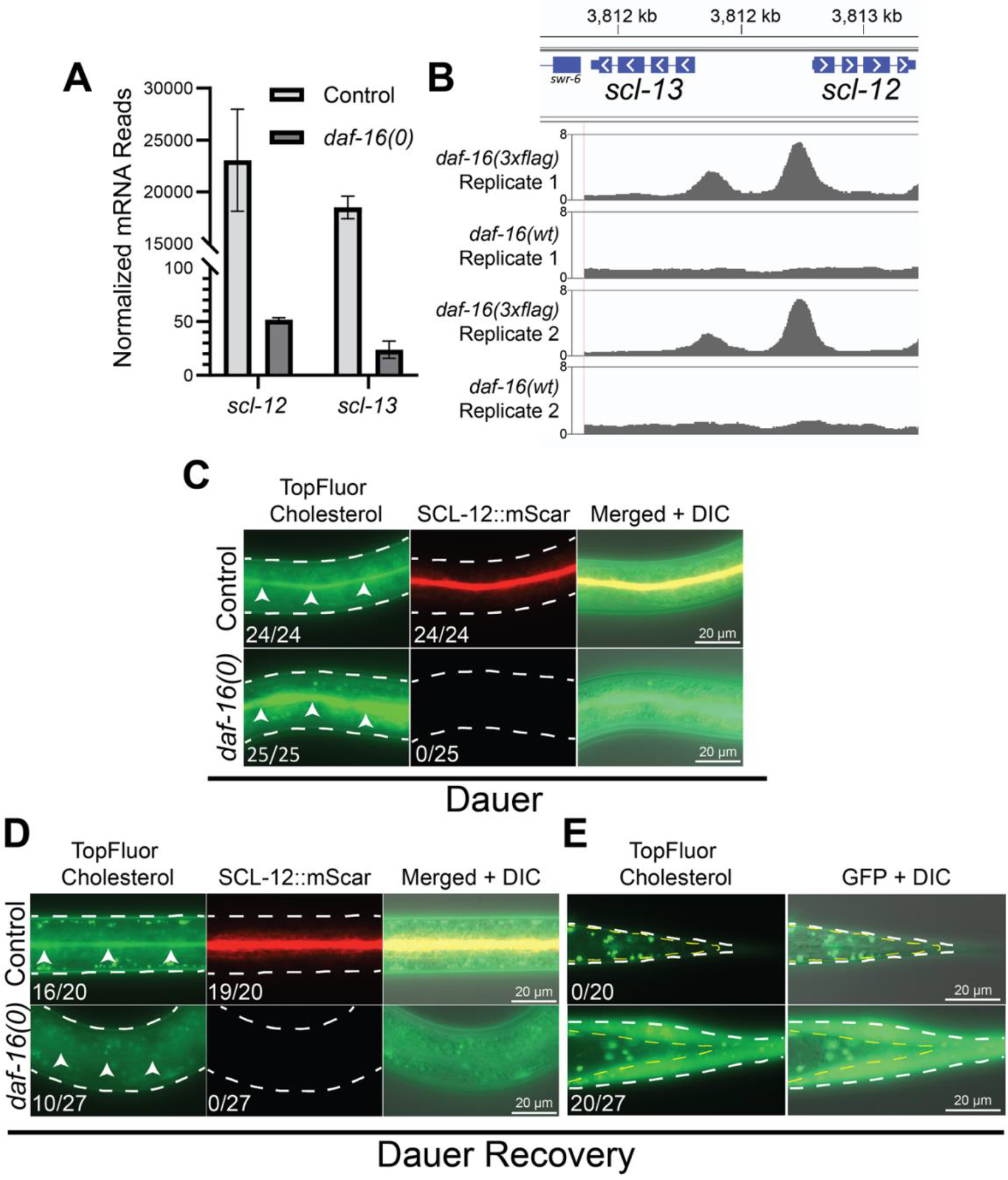
*daf-16* promotes the retention of intestinal cholesterol during dauer recovery. (A) *daf-16* promotes the expression of *scl-12* and *scl-13*. Normalized mRNA reads in control and *daf-16(0)* dauer larvae (FDR = 1.18 x 10^-124^ for both genes), from Wirick et al., 2021^5^. Error bars denote the standard deviation between two biological replicates. (B) DAF-16 binds upstream of *scl-12* and *scl-13*. DAF-16::3xflag binding peaks in two *daf-16(3xflag)* and two *daf-16(+)* (N2) replicates. Kilobases represent the relative location on chromosome V. Arrowheads represent the direction of transcription. Binding peaks and genomic locations were screen-captured from Integrative Genome View (Version 2.18.4). Data from Galagali et al. 2026^6^. (C) *daf-16(0)* dauer larvae sequestered TopFluor cholesterol but lack SCL-12::mScarlet expression in the intestinal lumen. (D) *daf-16(0)* larvae recovering from dauer lacked TopFluor cholesterol and SCL-12::mScarlet expression in the intestinal lumen. (E) TopFluor cholesterol in *daf-16(0)* larvae recovering from dauer localized at the posterior end of the larva, between the epidermis and the cuticle. (D-E) Dauer recovery was induced with 100nM Δ7-DA. Images were taken 7 hours after administration of Δ7-DA. (C-E) White dashed lines outline worm cuticles. In (E), there was a gap between the worm body and a cuticle; the worm body is indicated with dashed yellow lines. Numbers represent the number of larvae with detectable expression of TopFluor cholesterol or SCL-12::mScarlet over total number of larvae assessed. (C-D) Arrowheads point to the intestinal lumen.

### *daf-16* promotes cholesterol retention during dauer recovery

To address whether *daf-16* might regulate cholesterol sequestration during dauer we administered TopFluor™ Cholesterol (Avanti Research), which reports cholesterol localization, to larvae during dauer formation and saw that cholesterol was indeed sequestered in the intestinal lumen in both control and *daf-16(0)* dauer larvae (Figure 3C). We also observed the expression of the endogenously tagged *scl-12* allele, *scl-12::mScarlet*^22^, in the strains assessed for TopFluor Cholesterol. In agreement with our mRNA-sequencing data, SCL-12::mScarlet expression was not detectable in *daf-16(0)* dauer larvae (Figure 3C).

As wild-type larvae recover from dauer, cholesterol bound to SCL-12 is mobilized from the gut lumen into intestinal cells and released to become available for DA biosynthesis^22^. We asked if cholesterol was properly trafficked as *daf-16(0)* dauer larvae were recovering. To drive dauer recovery, we treated control and *daf-16(0)* dauer larvae, that were previously fed TopFluor Cholesterol, with 100nM Δ7-DA. After 7 hours of incubation on plates containing Δ7-DA, we noticed that a large fraction of *daf-16(0)* larvae no longer contained detectable TopFluor Cholesterol signal in the gut lumen, and instead the signal accumulated in what appeared to be an expanded cuticle region at the posterior end of the larvae (Figure 3D-E). In contrast, TopFluor Cholesterol was still visible in the intestinal lumen of control larvae at this same time point (Figure 3D). SCL-12::mScarlet remained undetectable in *daf-16(0)* larvae during dauer recovery (Figure 3D). These findings indicate that *daf-16(0)* larvae fail to retain cholesterol in the gut lumen upon dauer recovery, resulting in the loss of cholesterol that was stored during dauer, possibly by defecation.

The failure to retain cholesterol observed in recovering *daf-16(0)* dauer larvae is a related but distinct phenotype from the failure to sequester cholesterol observed in *scl-12(0) scl-13(0)* dauer larvae^22^. Residual expression of *scl-12* and *scl-13* in *daf-16(0)* dauer larvae could be sufficient to sequester cholesterol during dauer, but not to retain cholesterol during dauer recovery. This possibility seems unlikely because expression of *scl-13* in the intestine of pre-dauer and dauer larvae was not sufficient to suppress *daf-16(0)* phenotypes (Figure S7). Alternatively, there may be other compensatory mechanisms that can sequester, but not retain, cholesterol, such as the 4 additional *scl* genes (*scl-1*, *scl-2*, *scl-11*, and *scl-21*) that are misregulated in *daf-16(0)* dauer larvae^5^.

### *daf-16* functions prior to dauer formation to regulate post-dauer adult cell fate

SCL-12::mScarlet expression begins prior to dauer entry as cholesterol sequestration initiates^22^. If *daf-16* promotes PD adult cell fate by activating expression of *scl-12* and related genes, thereby enabling cholesterol retention and DA production, then we reasoned that *daf-16* would be required before dauer formation to specify adult cell fate in PD adults. In contrast, if *daf-16* acts more directly in DA biosynthesis or in transcription of *let-7-*family miRNAs, then *daf-16* would be required during or after dauer. To test if *daf-16* regulates adult cell fate before or after dauer formation, we used the AID system^23^ to conditionally deplete DAF-16::AID throughout somatic tissue either before dauer formation, after dauer formation had occurred, or both before and after dauer formation, and assessed alae formation in PD adults (Figure 4A). We found that depletion of DAF-16::AID before dauer resulted in alae defects, regardless of whether DAF-16::AID was depleted after dauer (Figure 4B). In contrast, depletion of DAF-16::AID only after dauer did not produce alae defects (Figure 4B).

**Figure 4.**
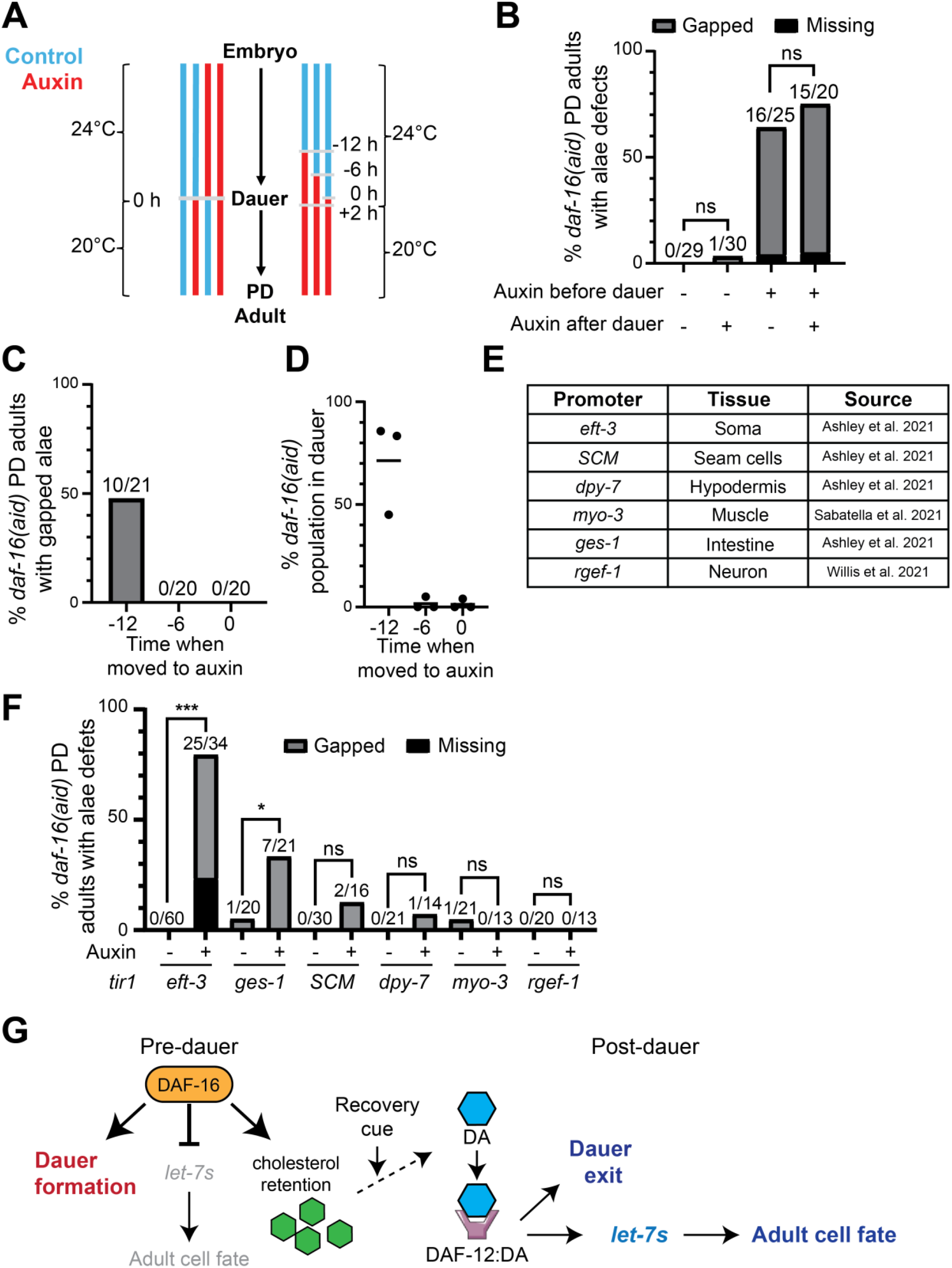
DAF-16 functions before dauer in the intestine to promote adult cell fate after dauer. (A) Model for auxin treatments in B-D, where time 0 = the approximate time when larvae entered dauer (48 hours after embryo synchronization). The left side refers to data shown in panel B, and the right refers to data shown in panels C-D, where dauer larvae were shifted from 24°C to 20°C approximately 2 hours after dauer entry (50 hours after embryo synchronization). Blue bars indicate times that larvae were on NGM plates and red bars indicate times on auxin plates. Gray bars represent time points that larvae were passed to a new plate. (B) Ubiquitous depletion of DAF-16 before dauer resulted in defects in adult alae after dauer. (C-D) DAF-16 is required between 12 and 6 hours before dauer formation to promote the expression of adult alae after dauer. Depletion of DAF-16 12 hours before dauer entry resulted in defects in adult alae after dauer (C) and dauer recovery (D). (D) Each data point represents the population of animals remaining in dauer 24 hours after shifting dauer larvae from 24°C to 20°C in an individual experiment (n=18-25 larvae per data point). (E) Table of tissue specific *tir1* drivers used in panel F^37–39^. (F) Depletion of DAF-16 in intestinal tissue resulted in adult alae defects after dauer. ***p<0.001, *p=0.044, 2X3 Fisher’s exact test, two-tailed. (G) Model for the coordination of dauer entry and exit with the regulation of stage-specific seam cell fate by *daf-16*.

To narrow down the window when *daf-16* is required, we depleted DAF-16::AID for different amounts of time before dauer formation as well as during dauer formation. Specifically, we began auxin treatment at 12, 6, or 0 hours before dauer formation. Approximately 2 hours after dauer formation, larvae were shifted to 20°C to induce dauer recovery. Alae were later assessed in worms that had developed into PD adults (Figure 4A). We found that depletion of DAF-16::AID during dauer or 6 hours before dauer formation did not result in PD alae defects, while depletion 12 hours before dauer formation did produce alae defects (Figure 4C). We also assessed dauer exit for these same conditions and found similar results, where only depletion of DAF-16::AID 12 hours prior to dauer formation produced a dauer exit phenotype (Figure 4D). Taken together, these results indicate that DAF-16 activity is required starting before dauer to promote adult seam cell fate in PD adults.

### *daf-16* functions cell non-autonomously to regulate post-dauer adult cell fate

If *daf-16* regulates dauer recovery and PD adult cell fate through promoting the retention of cholesterol upon dauer recovery, then we predicted that *daf-16* would be required in the intestine for these processes. We again used the AID system to deplete DAF-16::AID throughout the dauer life history but expressed TIR1 under various tissue-specific promoters (Figure 4E). While broad somatic depletion of DAF-16::AID produced the strongest phenotypes, we found that depletion of DAF-16::AID in the intestine resulted in a moderate, yet statistically significant, alae defect (Figure 4F). Furthermore, depletion DAF-16::AID in other tissues, including the hypodermis, caused low-penetrance alae defects that were not significantly different than the no-auxin control (Figure 4F). In contrast, none of the tissue-specific depletions recapitulated the dauer exit phenotypes produced by depletion of DAF-16::AID throughout the soma (Figure S8).

Similar to our observations using the *daf-16* null allele (Figure S4), we found that tissue-specific depletion of DAF-16::AID resulted in alae defects only in animals that were delayed in dauer recovery, consistent with the hypothesis that dauer recovery and adult cell fate are connected in this context (Figure S9). A possible explanation for this connection is that the regulation of cholesterol availability coordinates control of dauer exit and adult cell fate by modulating DAF-12 activity. In *daf-16(0)* recovering dauer larvae, we found that most, but not all, animals failed to retain TopFluor cholesterol in the gut lumen (Figure 3D). We speculate that *daf-16(-)* larvae that retain cholesterol during dauer recovery may produce enough DA to allow the activating DAF-12:DA complex to promote dauer recovery without delay and to allow the expression of adult cell fate in PD adults.

### *daf-16* coordinates dauer entry with the anticipation of dauer exit and post-dauer development via cholesterol retention and DAF-12:DA activity

Our findings that *daf-16* opposes adult cell fate during dauer and promotes adult cell fate after dauer suggest that *daf-16* is a key factor that coordinates the decision to enter and remain in dauer diapause with the developmental consequences of dauer arrest. Combining our findings with prior work on *daf-16,* we propose the following model (Figure 4G). During the predauer L2d stage, when IIS activity is low, DAF-16 is active and promotes entry into dauer diapause^2^. In addition, *daf-16* opposes the premature expression of *let-7-*family microRNAs and adult enriched collagens, preserving larval cell fate during developmental arrest^5,6^. Remarkably, at the same time as *daf-16* is acting to promote dauer formation, *daf-16* is also setting the stage to allow dauer exit and post-dauer developmental progression via regulation of DAF-12 ligand. Specifically, we propose that *daf-16* acts before dauer formation to promote the retention of cholesterol in the gut lumen upon dauer recovery. This cholesterol will then be available to serve as a substrate for the biosynthesis of DA which would then bind to DAF-12. The activating DAF-12:DA complex can then promote dauer exit. Finally, in PD larvae, DAF-12:DA can promote the transcription of the *let-7* family miRNAs in PD larvae, leading to the transition from larval to adult seam cell fate.

Our work provides new insight into the mechanisms by which a developmental pathway is modulated to accommodate a variable and potentially lengthy developmental arrest. The central role for *daf-16* in this process is intriguing given the conserved function for FOXO proteins in promoting diapause across species^24,25^. Since the role for *let-7-*family miRNAs in promoting differentiation is also conserved in mammals and other species^26^ it will be interesting to determine the connection between FOXO proteins and *let-7* in other systems.

## Methods

### Strains and maintenance

All *C. elegans* strains were grown at 15°C or 20°C on standard Nematode Growth Medium (NGM) plates and seeded with *E. coli* OP50^27^. Strains used in this study can be found in Table S1.

### Synchronizing worms in continuous development

For Figures 1D-E, gravid adults were allowed to lay embryos at 25°C for 6 hours. Adults were removed from the plates and embryos were allowed to develop at 25°C for 24 hours, then were shifted to 15°C until they reached adulthood to match conditions under which larvae were grown in the dauer life history.

### Dauer induction

All strains that developed through the dauer life history in this study contained the *daf-7(e1372)* temperature sensitive allele which forms dauer larvae at 24-25°C^3,28^. With exceptions noted below, populations of embryos were obtained from plates with gravid adults that were treated with two 2-minute incubations of 1M NaOH, 0.5% sodium hypochlorite, then washed twice with dH_2_O. Embryos were plated onto NGM plates seeded with OP50 and incubated at 24°C for approximately 50 hours to allow larvae to enter dauer.

For Figures 1C-E and S1, gravid adults were allowed to lay embryos on NGM plates seeded with OP50 for 2-8 hours at 24-25°C. Adults were removed from the plates and embryos were incubated at 24-25°C for 48-52 hours to allow dauer larvae to form.

### Post-dauer populations

Once populations of dauer larvae were obtained, they were picked to new NGM plates seeded with OP50 and shifted to 20°C, unless otherwise specified. For Figures 1C-E, dauer larvae were recovered at 15°C.

Generally, *daf-16(+)* PD adults were assessed 2 days after shifting dauer larvae from 24°C to 15-20°C. Larvae that exited dauer and reached adulthood on the second day were assessed on day 2 for adult alae. To obtain day 3 PD adults, the adults that were not assessed on day 2 were picked off the plates, and the remaining larvae were incubated at 20°C for an additional day (day 3) before assessment.

### Compound microscopy

Animals were mounted onto glass slides with 2% agarose pads and immobilized with 0.1M levamisole. A Zeiss AxioImager D2 compound microscope with HPC 200 C or X-cite Xylis II fluorescent optics was used to image animals. DIC and fluorescence images were captured with a AxioCam MRm Rev 3 or Axiocam 807 mono camera and ZEN 3.2 or 10 software. GFP images were captured with a high efficiency GFP shift free filter, mScarlet images were captured with a Zeiss 63HE filter, and YFP images were captured with a Zeiss 46HE filter. To adjust for TopFluor Cholesterol signal output, the GFP brightness display range was set to 0-2600 for Figure 3C-D and 0-5000 for Figure 3E on ZEN 10 software.

### Phenotypic analysis

<u>Adult alae</u> in adults were observed with DIC optics along the lateral cuticle of the worm, between the rectum and pharynx. An adult was classified to have “gapped” alae if alae were present above some but not all seam cells, while worms lacking any detectable alae were classified as “missing”. <u>Dauer exit:</u> Populations of dauer larvae were obtained as stated above. Approximately 30-50 dauer larvae were picked to 60mm NGM plates seeded with OP50 and incubated at 20°C for 24 hours. Dauer larvae were distinguished from recovering dauer larvae by radial constriction and pharyngeal pumping. The number of both dauer and non-dauer larvae present on the NGM were counted to give a percentage of larvae remaining in dauer. Larvae that crawled up the side of the petri dish were disregarded.

### Fluorescent reporters

For *col-19p::gfp* expression in Figure 1C, an animal was determined to express *col-19p::gfp* if any hypodermal nuclei had detectable expression.

The *<u>ajm-1::gfp</u>* reporter was used to determine the presence of adherens junctions. Worms were classified with having junctions when the expression of *ajm-1::gfp* was detectable between seam cells.

To determine the penetrance of *<u>col-38p::yfp</u>*, L2d larvae were assessed 28hr after embryo laying, L2d molt larvae were assessed 38 hours after embryo laying, and dauer larvae were assessed 48hr after embryo laying. For post-dauer stages, dauer larvae were picked to fresh plates to recover at 20°C. PDL3 and PDL4 larvae were assessed for *col-38p::yfp* expression 24-72hr after placement at 20°C. To verify larvae were not yet in dauer (L2d) or had recovered from dauer (PDL3), fluorescent beads were used to demonstrate that larvae were pumping and consuming food^15^. The extension of gonad arms was used to distinguish between PDL3 and PDL4 larvae. Adults were determined by the presence of embryos.

Fluorescence of *<u>let-7p::gfp</u>* was measured using ImageJ (Version 1.54f) software. Images of an entire larva were captured with DIC and GFP channels on a 10X (Figure 1G) or 20X (Figure 2D) objective. The entire body of a larva from a DIC image was selected as the region of interest (ROI) using the freehand selection tool. That ROI was then measured on the GFP image to give the mean fluorescence intensity of GFP signal for a given worm.

The presence of <u>TopFluor™ Cholesterol</u> (Avanti Research, CAS: 878557-19-8) sequestration and <u>SCL-12::mScarlet</u> expression in the gut lumen were assessed in dauer larvae and recovering dauer larvae. Recovering dauer larvae were also assessed for TopFluor Cholesterol outside of the worm body. The expression of TopFluor Cholesterol in the gut lumen was assessed by examining the portion of the intestine directly posterior to the pharynx.

### PDL4 collection for Taqman qPCR

Populations of PDL4 larvae were obtained as described above. PDL4 larvae were picked into 100µL of dH_2_O in tubes, then 1mL TRI-Reagent (Ambion) was added. Samples were incubated at room temperature for 10 minutes before flash freezing. Samples were stored at -80°C.

### RNA Isolation

Total RNA isolation was conducted using TRI-Reagent (Ambion) standard protocol with the following modifications: to improve extraction efficiency pelleted *C. elegans* in TRI-Reagent were subjected to three freeze/thaw/vortex cycles prior to BCP addition, isopropanol precipitation was conducted in the presence of glycogen for 2 hours at -80°C, RNA was pelleted by centrifugation at 4°C for 30 minutes at 20,000 x g; the pellet was washed three times in 70% ethanol and resuspended in water. Thermo Scientific Nanodrop2000 was used to determine the concentration and quality of the RNA prior to TaqMan synthesis.

### Taqman qPCR analysis

MiRNA levels were quantified with a CFX96 Real-Time System (BioRad) using TaqMan assays and probes (Applied Biosystems hsa-let-7a (assay# 000377), cel-miR-48 (assay# 000208), cel-miR-84 (assay# 463262_mat), cel-miR-241(assay# 000249), and U18 (assay# 001764) according to the vendor protocol with modifications as described in Billi et al. 2012^29^. Relative RNA levels were calculated based on the ΔΔ2Ct method^30^ using U18 for normalization. Results presented are the average of independent calculations from biological duplicates.

### Conditional depletion of proteins

Strains containing the *daf-16(ot975)* or *daf-12(ot874)* were treated with a final concentration of 4mM Naphthaleneacetic Acid (KNAA) (PhytoTech Labs, CAS: 15165-79-4) in dH_2_O dissolved in NGM media. To correct for the volume of KNAA solution added to the molten NGM, the volume of KNAA to be added was removed from the total volume of water in the NGM media prior to adding the NGM ingredients.

For Figure 4B, 4F, Figures S8-9, embryos were obtained by sodium hypochlorite treatment and put onto 4mM KNAA plates or NGM plates seeded with 10X concentrated OP50 and grown at 24°C to induce dauer formation. Dauer larvae were picked to new 4mM KNAA plates or NGM plates seeded with 10X concentrated OP50 and recovered from dauer at 20°C.

For Figures 4C-D, embryos were obtained by sodium hypochlorite treatment and put onto NGM plates seeded with OP50 and incubated at 24°C. At 36, 42 and 48 hours of incubation at 24°C on NGM (12, 6, and 0 h before dauer formation), larvae were washed off of the plates with 1M M9 buffer, centrifuged, washed with M9 buffer, distributed to 4mM KNAA plates seeded with 10X concentrated OP50, and incubated at 24°C. Once 50 hours of total incubation at 24°C elapsed (2 h after dauer formation), dauer larvae were picked to new 4mM KNAA plates seeded with 10X concentrated OP50 and recovered from dauer at 20°C. Dauer recovery was assessed in a similar fashion as stated above, but on NGM plates with/without KNAA present.

For Figure S5, dauer larvae were obtained were obtained by sodium hypochlorite treatment added to NGM plates seeded with OP50 and incubated at 24°C. Dauer larvae were recovered on NGM plates seeded with OP50 at 20°C for 12 hours before being transferred to NGM or 4mM KNAA plates seeded with 10X concentrated OP50. Larvae were then incubated at 20°C for an additional 1.5 days before assessing adults.

### Daf-c assessment of *daf-12(ot874)*

Synchronized populations of embryos were obtained by sodium hypochlorite treatment and placed onto NGM plates seeded with OP50. Plates with embryos were incubated at 20°C for 3 days, then the number of dauer and non-dauer animals were counted. Dauer larvae were distinguished from non-dauer animals by radial constriction and pharyngeal pumping.

### Exogenous sterol treatments

Cholesterol (Sigma Aldrich, CAS: 57-88-5), 7-Dehydrocholesterol (7DHC) (Cayman Chemical, CAS: 434-16-2), Lathosterol (Cayman Chemical, CAS: 80-99-9), and Cholestenone (Cayman Chemical, CAS: 601-57-0) were suspended in 200 proof ethanol to a concentration of 33.3mM. (25S)-Δ^7^-Dafachonic acid (DA) (Cayman Chemical, CAS: 949004-12-0) was suspended in 200 proof ethanol to a concentration of 100µM. To test the effect of these sterol molecules, dauer larvae were picked to NGM plates seeded with 10X concentrated OP50 with a single sterol/ethanol solution to a final concentration of 33.3µM for cholesterol, 7DHC, Lathosterol, and Cholestenone, or 100nM for Δ7-DA, based on the volume of the NGM plate. Dauer larvae were then recovered at 20°C.

To determine the best concentration of Δ7-DA to use in this study, we treated *daf-16(0)* dauer larvae with serial dilutions of Δ7-DA. We found that 100nM promotes most *daf-16(0)* dauer larvae to exit dauer (Figure S10).

### TopFluor™ Cholesterol treatment

TopFluor Cholesterol was dissolved in 200 proof ethanol to a concentration of 2.5mg/mL. NGM plates without cholesterol added were seeded with 10X concentrated OP50 and TopFluor Cholesterol (final concentration of TopFluor Cholesterol:media, 1:1000). Synchronized populations of embryos were obtained by sodium hypochlorite treatment (see above) and placed onto TopFluor Cholesterol plates. To drive dauer recovery, dauer larvae were recovered on NGM plates containing 100nM Δ7-DA seeded with 10X concentrated OP50.

### Transgene construction

The *ets-10* promoter included the sequence from the 5’ primer given in Shih et al. 2019^31^ to the start codon. The *scl-13* sequence was derived from WormBase (WS297). The spliced sequence was used, but a 5’ synthetic intron was added at the location where the first intron of *scl-13* is found. A linker and 3xflag sequence were added immediately upstream of the stop codon. The *unc-54* 3’UTR sequence was as described in Shih et al. 2019^31^. The mCherry construct was identical to the *scl-13* construct except the mCherry coding sequence was taken from Crane et al. 2019^32^ and included the same synthetic intron as used for *scl-13.* The constructs were ordered as Gene Fragments from Twist Bioscience. Relevant sequence information is found below.

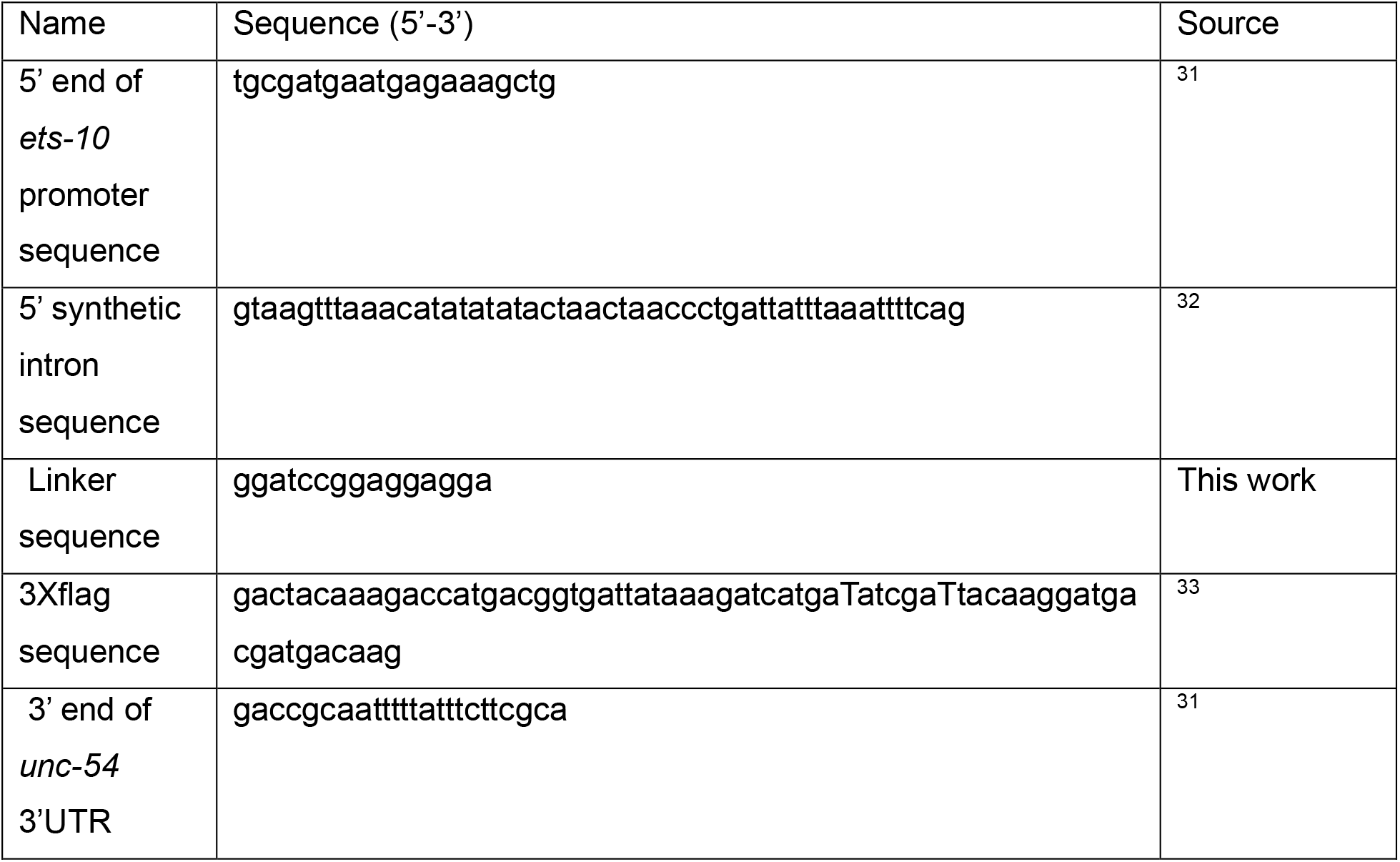

### Transgenic strains

To create *C. elegans* strains that express *scl-13* or *mCherry* under the *ets-10* promoter in extrachromosomal arrays, *daf-16(mgDf50); daf-7(e1372) pha-1(e2123)* gravid adults were injected with a mix containing 20 ng/µL *scl-13* or *mCherry* construct, 20 ng/µL *ceh-22::gfp*, 20 ng/µL pBX *(pha-1(+))*, and 40 ng/µL L4440. Injected gravid adults were singled out onto NGM plates seeded with OP50 and incubated at 20°C. F1 larvae were selected based on *pha-1* rescue and expression of *ceh-22::gfp*. F2 populations that retained the array were maintained as individual lines.

### Reproducibility, graphs, and statistical analysis

All data presented were collected from at least two independent experiments performed on different days. Graphs were generated with GraphPad Prism (Version 10.4.1). All statistical analyses were conducted on GraphPad Prism (Version 10.4.1) with the exception of Fisher’s Exact Tests which were conducted on VassarStats (vassarstats.net). Statistical analyses that provided a p-value <0.05 were deemed significant. Asterisks denoting significance were used as follows: *p<0.05, **p<0.01, and ***p<0.001. Statistical tests used for each experiment can be found in the respective figure legend.

## Acknowledgements

We thank Luisa Cochella (Johns Hopkins University) and Oliver Hobert (Columbia University) for sharing unpublished reagents. We thank Iva Greenwald (Columbia University), Kathrin Schmeisser (MPI-CBG), and Karp lab members for useful discussions. We thank Nehul Tanna (Karp lab) for injecting worms. Some strains were provided by the CGC, which is funded by NIH Office of Research Infrastructure Programs (P40 OD010440). We thank WormBase and the Alliance of Genome Resources^34^. This work was supported by NSF CAREER 1352283 (to XK), NIH R15GM150082 (to XK).

