## Supplemental Figures and Tables for "*daf-16/FOXO* activity before dauer diapause anticipates dauer exit and post-dauer development via DAF-12/NHR"

### Supplemental Materials

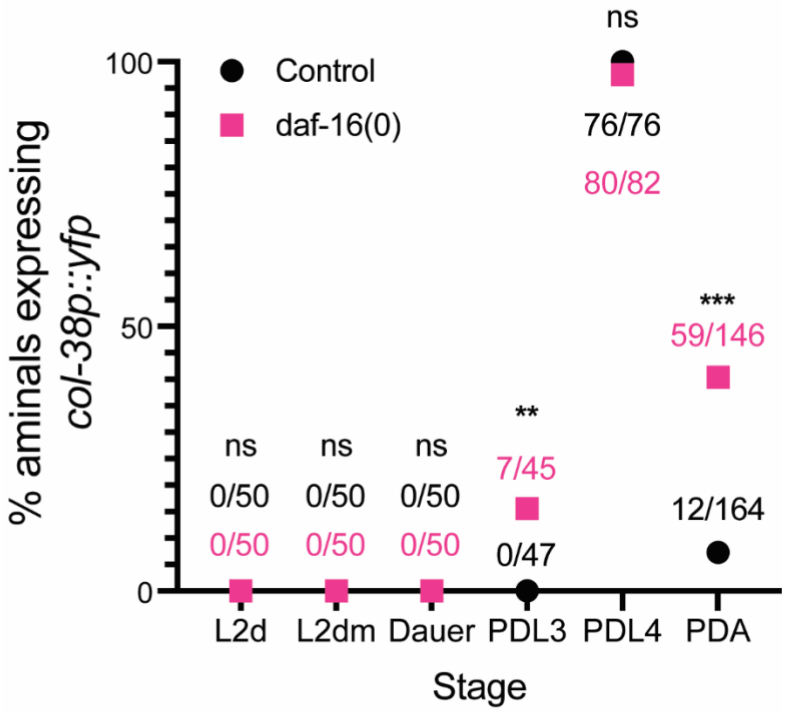


**Figure S1. *daf-16(0)* PD adults failed to properly downregulate *col-38p::yfp* in adulthood.** *daf-16(0)* mutants expressed the L4 marker *col-38p::yfp*[1] in PDL4, but failed to properly downregulate this marker in adulthood. Numbers indicate the number of animals expressing *col-38p::yfp* over the total number of animals assessed. **p=0.005, ***p<0.001, 2X2 Fisher's exact test, two tailed.

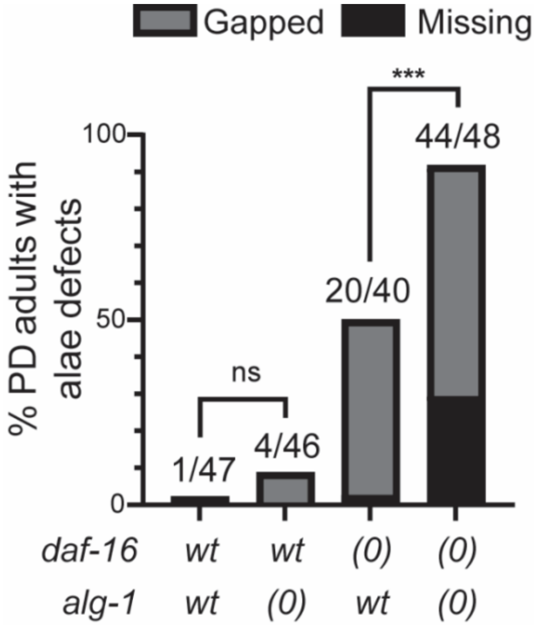


**Figure S2. *daf-16(0)* and *alg-1(0)* alleles showed mutual enhancement for defects in adult alae after dauer**. Numbers indicate the number of animals with either gapped or missing alae over the total number of animals assessed. *wt* = wild-type allele; *(0)* = null mutant allele. ***p<0.001, 2X3 Fisher's exact test, two-tailed.

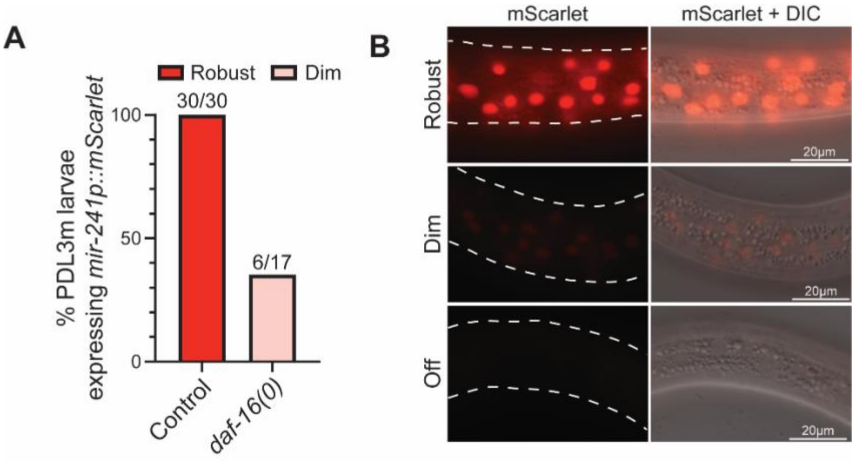


**Figure S3. *daf-16* promotes *mir-241p::mScarlet* expression after dauer.** (A) Control PDL3m larvae robustly expressed *mir-241p::mScarlet* while *daf-16(0)* PDL3m larvae expressed this reporter dimly and at low penetrance. Numbers indicate the number of larvae expressing any detectable level of *mir-241p::mScarlet* over the total number of larvae observed. (B) Representative images PDL3m larvae expressing robust, dim, and no *mir-241p::mScarlet*. Worm bodies are outlined in white dashed lines. All fluorescence images were taken at identical settings.

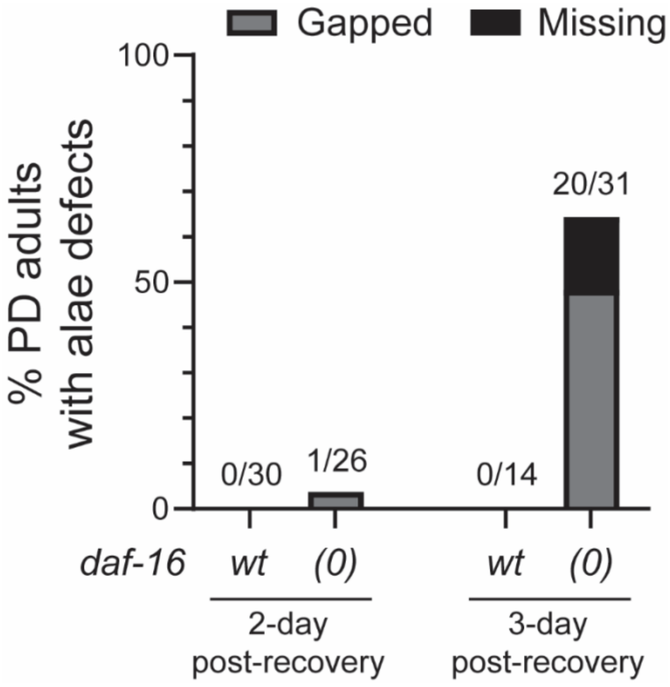


**Figure S4. The dauer recovery defects in *daf-16(0)* mutants correlate with defects in adult alae.** Control *daf-16(+)* PD adults did not display defects in adult alae even in worms with delayed dauer recovery. *daf-16(0)* PD adults that recover from dauer and reach adulthood in 2 days had substantially fewer defects in adult alae than those that reached adulthood 3 days after recovery.

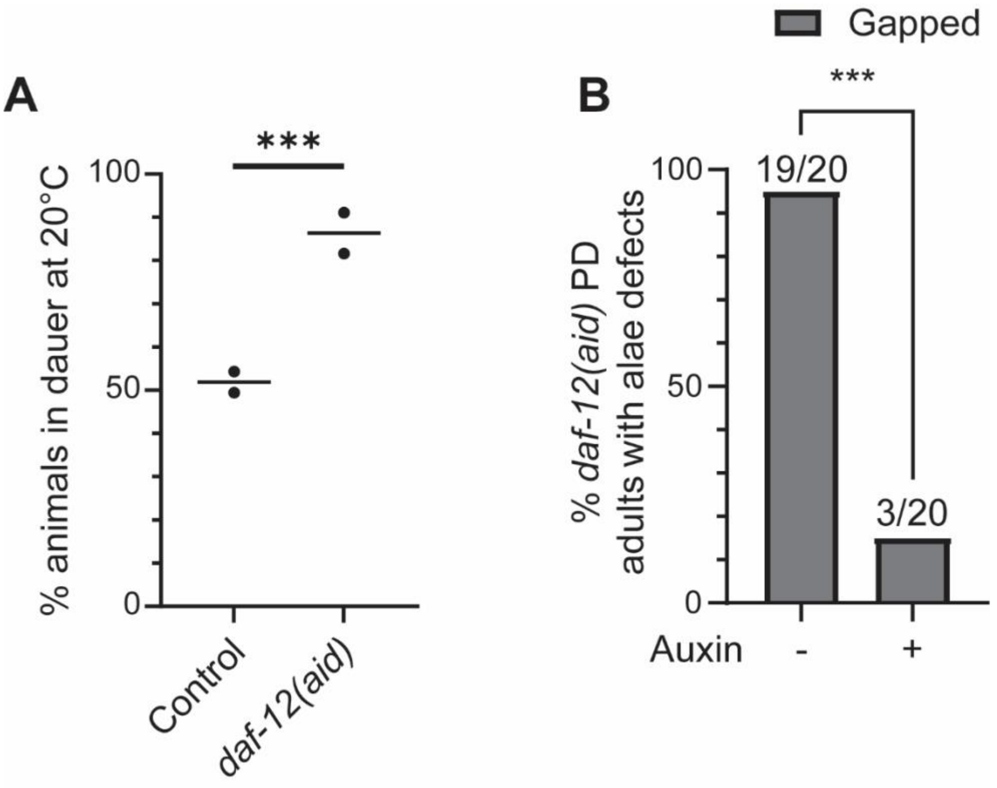


**Figure S5. The *daf-12(ot875[daf-12::TagRFP::aid])* allele shows dauer-constitutive and alae phenotypes similar to *daf-12* ligand-binding mutants.** (A) The *daf-12(aid)* strain entered dauer at 20°C more readily than the control *daf-7(e1372)* strain. ***p<0.001, Pearson's Chi Squared test. Each data point represents an individual population that was synchronized as embryos and grown at 20°C for 3 days (n=167-962 larvae per data point). Black bars represent the mean % of larvae in dauer. Note that these strains lack *tir-1* drivers. (B) PD adults harboring the *daf-12(aid)* allele had penetrant defects in alae formation which were suppressed when DAF-12 was depleted. ***p<0.001, 2X2 Fisher's exact test, one-tailed. Dauer formation and then recovery was induced in the *eft-3p::tir1; daf-7(e1372); daf-12(aid)* strain before addition of auxin. Dauer larvae were recovered at 20°C for 12 hours on NGM plates before being transferred to auxin plates until adulthood. Since *daf-12* is required for dauer formation[2], we originally attempted to use the AID system to conditionally deplete *daf-12* in post-dauer animals. In favorable conditions, the DAF-12:DA complex activates *let-7*-family miRNA transcription and promotes continuous development. In adverse conditions, ligand-free DAF-12 binds its corepressor DIN-1S to repress *let-7*-family miRNA transcription and promote dauer entry[2–5]. *daf-9(0)* mutants that fail to produce DA, or *daf-12(rh273)* mutants that bind DA poorly, shift the DAF-12 complex to its repressive form and display dauer-constitutive and reiterative phenotypes[2,6]. We speculate that the ligand-binding site is slightly less available in the DAF-12 protein produced by the *daf-12(aid)* allele compared to wild-type DAF-12, leading to the dauer exit and PD adult cell fate defects seen here.

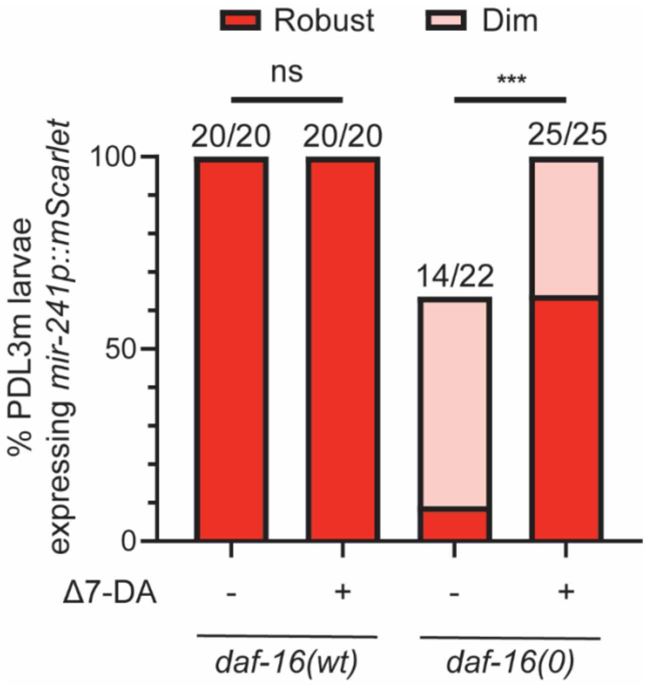


**Figure S6. Treatment with Δ7-DA rescued *mir-241p::mScarlet* expression in *daf-16(0)* dauer larvae.** All *daf-16(0)* PDL3 molt larvae expressed *mir-241p::mScarlet* when treated with Δ7-DA, while ethanol treated mutants did not. For representative images of categories, see Figure S3B.

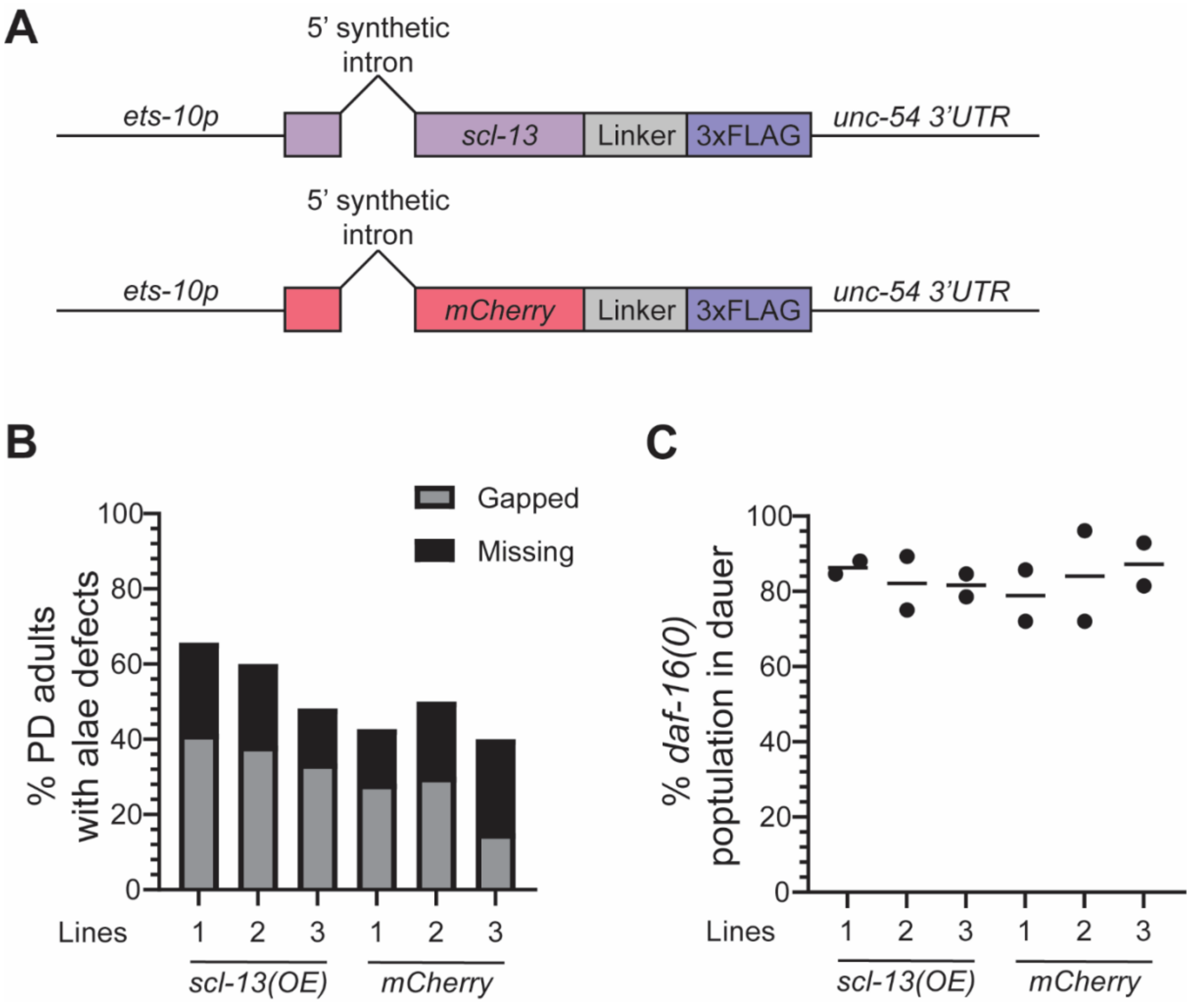


**Figure S7. Intestinal expression of *scl-13* did not suppress *daf-16(0)* phenotypes.** (A) Transgene models for intestinal expression of *scl-13* (top) and mCherry (bottom). *scl-13* and mCherry cDNAs were placed under the *ets-10* promoter with a 5' synthetic intron after the first exon[7,8]. Numbers 1-3 indicate three independent lines. Defects in adult alae (B) and dauer exit (C) were not suppressed in *daf-16(0)* animals harboring any of the three *scl-13* extrachromosomal arrays.

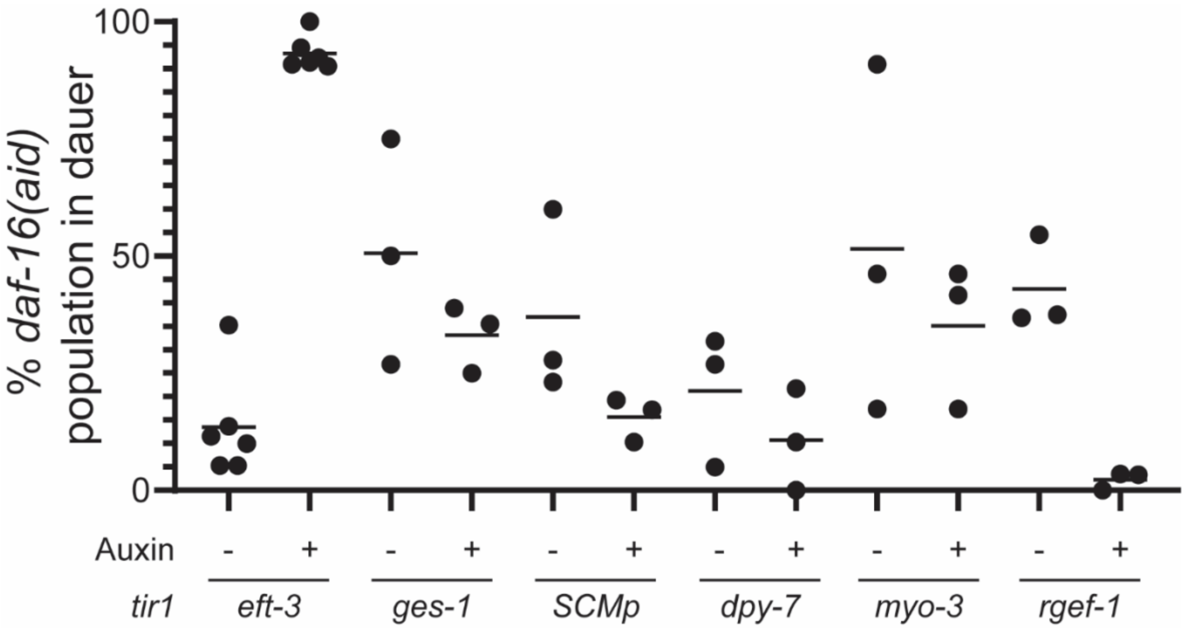


**Figure S8. Tissue-specific depletion of DAF-16 did not compromise the timing of dauer exit.** Depletion of DAF-16 throughout somatic tissue recapitulated the dauer exit defects in *daf-16(0)* mutants (Figure 1F), but depletion from intestine, seam cells, hypodermis, body-wall muscle, or neurons did not cause a clear difference from no-auxin controls. Most tissue-specific *tir1*-expressing strains displayed some background dauer exit defects compared to the *eft-3p::tir1* strain. Each data point represents the population of larvae remaining in dauer 24 hours after shifting dauer larvae from 24°C to 20°C in an individual experiment (n=11-30 larvae per data point).

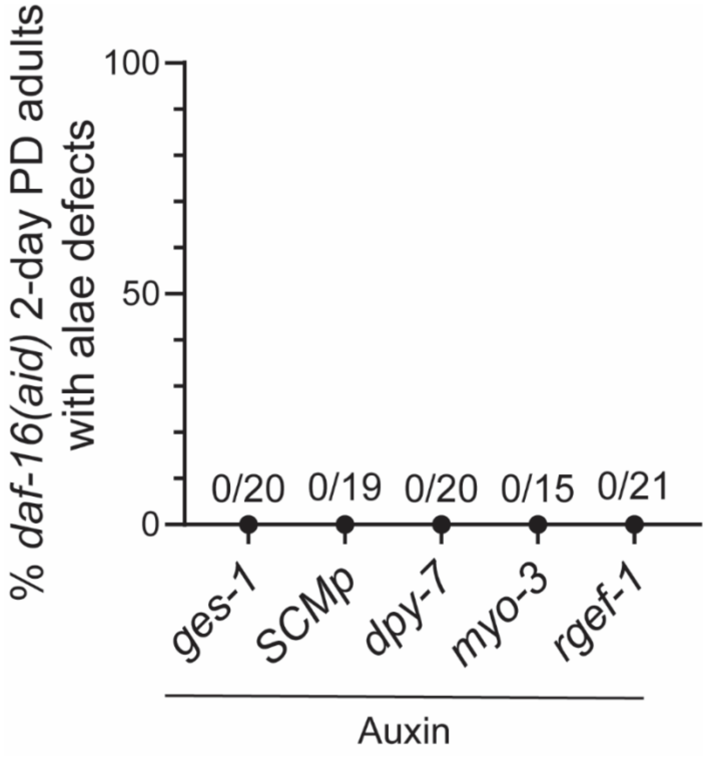


**Figure S9. Depletion of DAF-16 in different tissues did not affect adult alae formation in 2-day post-recovery adults.** Note that the *eft-3p::tir1* control is not included because none of these worms reached adulthood within 2 days after dauer recovery. The animals shown in Figure 4F reached adulthood 3 days after recovery from dauer diapause.

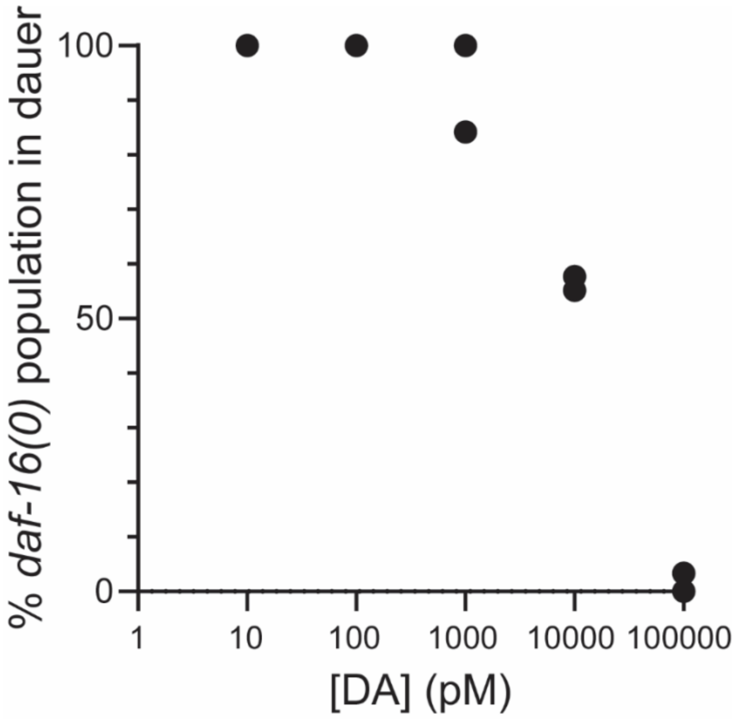


**Figure S10. Dose dependent curve for the concentration of dafachronic acid required to suppress dauer recovery defects in *daf-16(0)* dauer larvae.** Each data point represents the percentage of larvae remaining in dauer 24 hours after shifting dauer larvae from 24°C to 20°C, at a given concentration of dafachronic acid. At 100nM (100000 pM), most *daf-16(0)* larvae recovered from dauer in this time. n=19-30 larvae for each experiment. Two experiments are shown for each concentration; at 10 and 100 pM, 100% of the population remained in dauer.

**Table S1. List of strains used in this study.**

| Figure | Strain | Genotype | Transgene sources |
|---|---|---|---|
| 1C | VT1777 | *daf-7(e1372) III; maIs105[col-19p::gfp] V* | *maIs105*:[9] |
| | XV36 | *daf-16(mgDf50) I; daf-7(e1372) III; maIs105 V* | |
| 1D-E | XV24 | *wIs78[ajm-1::gfp + SCMp::gfp] IV* | *wIs78*:[10,11] |
| | XV27 | *daf-7(e1372) III; wIs78 IV* | |
| | XV28 | *daf-16(mgDf50) I; wIs78 IV* | |
| | XV29 | *daf-16(mgDf50) I; daf-7(e1372) III; wIs78 IV* | |
| 1F, 3A | CB1372 | *daf-7(e1372) III* | |
| | VT2317 | *daf-16(mgDf50) I; daf-7(e1372) III* | |
| 1G, 2D | QK152 | *xkIs43[let-7p::gfp] I; daf-7(e1372) III* | *xkIs43*:[12] |
| | XV285 | *daf-16(mgDf50) xkIs43 I; daf-7(e1372) III* | |
| 1H, 2E | XV319 | *daf-7(e1372) III; mir-241(umn47[mir-241p+SL1::egl-13-NLS::mScarlet-I::c-myc-NLS::linker::mODC(422-461)(E428A/E430A/E431A)::let-858 3' UTR]) V* | *umn47*: CGC |
| | XV320 | *daf-16(mgDf50) I; daf-7(e1372) III; mir-241(umn47) V* | |
| 2B-C | XV291 | *daf-16(mgDf50) I; daf-7(e1372) III; maIs105 V* | |
| 3B | GS8924 | *daf-16(ar620[daf-16::zf1::wrmScarlet::3xflag]) I* | |
| | N2 | Wild type | |
| 3C-E | XV325 | *daf-7(e1372) III; scl-12(trg1702[scl-12::mScarlet]) V* | *trg1702:*[13] |
| | XV326 | *daf-16(mgDf50) I; daf-7(e1372) III; scl-12(trg1702) V* | |
| 4B-D | XV281 | *daf-16(ot975[daf-16::mNeptune2.8::AID]) I; jsSi1579 cenSi1 [+ loxP eft-3p:TIR1::F2A::mTagBFP2::AID*::NLS::tbb-2 3'UTR *wrdSi57] (II:0.77); daf-7(e1372) III; maIs105 V* | *ot975*:[14]<br>*wrdSi57*:[15] |
| 4F | XV281 | *daf-16(ot975) I; jsSi1579 cenSi1 II; daf-7(e1372) III; maIs105 V* | |

| Figure | Strain | Genotype | Transgene sources |
| --- | --- | --- | --- |
| | XV334 | *daf-16(ot975) I; wrdSi44 II; daf-7(e1372) III; maIs105 V* | *wrdSi44:*[15] |
| | XV335 | *daf-16(ot975) I; wrdSi45 [dpy-7p::TIR1::F2A::mTagBFP2::AID*::NLS::tbb-2 3'UTR] (II:0.77); daf-7(e1372) III; maIs105 V* | *wrdSi45:*[15] |
| | XV336 | *daf-16(ot975) I; daf-7(e1372) III; emcSi71[myo-3p::TIR1::mRuby] (IV: -0.05); maIs105 V* | *emcSi71* [16] |
| | XV340 | *daf-16(ot975) I; reSi12 [ges-1p::TIR1::F2A::mTagBFP2::AID*::NLS::tbb-2 3'UTR] (II:0.77); daf-7(e1372) III; maIs105 V* | *reSi12:*[15] |
| | XV341 | *daf-16(ot975) I; kea15[rgef-1p::TIR1::mRuby::unc-54 3UTR + Cbr-unc-119(+)] II; daf-7(e1372) III; maIs105 V* | *kea15*[17] |
| S1 | XV128 | *daf-16(mgDf50) I; daf-7(e1372) III; deIs20 [col-38p::yfp]* | *deIs20*:[1] |
| | XV129 | *daf-7(e1372) III; deIs20* | |
| S2 | QK154 | *daf-16(mgDf50) I; daf-7(e1372) III; maIs105 V; alg-1(gk214) X* | |
| | VT1777 | *daf-7(e1372) III; maIs105 V* | |
| | XV88 | *daf-7(e1372) III; maIs105 V; alg-1(gk214) X* | |
| | XV291 | *daf-16(mgDf50) I; daf-7(e1372) III; maIs105 V* | |
| S3, S6 | XV319 | *daf-7(e1372) III; mir-241(umn47[mir-241p::mScarlet) V* | |
| | XV320 | *daf-16(mgDf50) I; daf-7(e1372) III; mir-241(umn47) V* | |
| S4 | VT1777 | *daf-7(e1372) III; maIs105[col-19p::gfp] V* | |
| | XV291 | *daf-16(mgDf50) I; daf-7(e1372) III; maIs105 V* | |
| S5A | CB1372 | *daf-7(e1372) III* | |
| S5A-B | XV337 | *jsSi1579 cenSi3 [+ loxP eft-3p:TIR1::F2A::mTagBFP2::AID*::NLS::tbb-2 3'UTR *wrdSi57] (II:0.77); daf-7(e1372) III; daf-12(ot874[daf-12::TagRFP::AID]) X* | |
| S7 | XV350 | *daf-16(mgDf50) I; daf-7(e1372) pha-1(e2123) III; cenEx4[ets-10p::scl-13(5' synthetic* | *cenEx4*: This study |

| Figure | Strain | Genotype | Transgene sources |
|---|---|---|---|
| | | *intron)::linker::3xFLAG::unc-54 3'UTR + ceh-22::gfp + pha-1(+)]* | |
| | XV351 | *daf-16(mgDf50) I; daf-7(e1372) pha-1(e2123) III; cenEx5[ets-10p::scl-13(5' synthetic intron)::linker::3xFLAG::unc-54 3'UTR + ceh-22::gfp + pha-1(+)]* | *cenEx5*: This study |
| | XV352 | *daf-16(mgDf50) I; daf-7(e1372) pha-1(e2123) III; cenEx6[ets-10p::scl-13(5' synthetic intron)::linker::3xFLAG::unc-54 3'UTR + ceh-22::gfp + pha-1(+)]* | *cenEx6*: This study |
| | XV353 | *daf-16(mgDf50) I; daf-7(e1372) pha-1(e2123) III; cenEx7[ets-10p::mCherry(5' synthetic intron)::linker::3xFLAG::unc-54 3'UTR + ceh-22::gfp + pha-1(+)]* | *cenEx7*: This study |
| | XV354 | *daf-16(mgDf50) I; daf-7(e1372) pha-1(e2123) III; cenEx8[ets-10p::mCherry(5' synthetic intron)::linker::3xFLAG::unc-54 3'UTR + ceh-22::gfp + pha-1(+)]* | *cenEx8*: This study |
| | XV355 | *daf-16(mgDf50) I; daf-7(e1372) pha-1(e2123) III; cenEx9[ets-10p::mCherry(5' synthetic intron)::linker::3xFLAG::unc-54 3'UTR + ceh-22::gfp + pha-1(+)]* | *cenEx9*: This study |
| S8-S9 | XV334 | *daf-16(ot975) I; wrdSi44 II; daf-7(e1372) III; maIs105 V* | |
| | XV335 | *daf-16(ot975) I; wrdSi45 [dpy-7p::TIR1::F2A::mTagBFP2::AID*::NLS::tbb-2 3'UTR] (II:0.77); daf-7(e1372) III; maIs105 V* | |
| | XV336 | *daf-16(ot975) I; daf-7(e1372) III; emcSi71[myo-3p::TIR1::mRuby] (IV: -0.05); maIs105 V* | |
| | XV340 | *daf-16(ot975) I; reSi12 [ges-1p::TIR1::F2A::mTagBFP2::AID*::NLS::tbb-2 3'UTR] (II:0.77); daf-7(e1372) III; maIs105 V* | |
| | XV341 | *daf-16(ot975) I; kea15[rgef-1p::TIR1::mRuby::unc-54 3UTR + Cbr-unc-119(+)] II; daf-7(e1372) III; maIs105 V* | |
| S10 | XV291 | *daf-16(mgDf50) I; daf-7(e1372) III; maIs105 V* | |
